# Impacts of host phylogeny, feeding styles, and parasite attachment site on isotopic discrimination in helminths infecting coral reef fish hosts

**DOI:** 10.1101/2020.05.31.126110

**Authors:** Philip M. Riekenberg, Marine J. Briand, Thibaud Moléana, Pierre Sasal, Marcel van der Meer, David W. Thieltges, Yves Letourneur

## Abstract

Stable isotopes of carbon and nitrogen characterize trophic relationships in predator-prey relationships, with clear differences between consumer and diet (discrimination factor, Δ^13^C, Δ^15^N). However, parasite-host isotopic relationships remain unclear, with Δ^13^C and Δ^15^N remaining incompletely characterized, especially for helminths. In this study, we used stable isotopes to determine discrimination factors for 13 parasite-host pairings of helminths in coral reef fish. Δ^15^N differences grouped according to phylogeny and attachment site on the hosts: Δ^15^N was positive for trematodes and nematodes from the digestive tract and varied for cestodes and nematodes from the general cavity. Δ^13^C showed more complex patterns with no effect of phylogeny or attachment site. A negative relationship was observed between Δ^15^N and host δ^15^N value among different host-parasite pairings as well as within 7 out of the 13 parings, indicating that host metabolic processing affects host-parasite discrimination values. In contrast, no relationships were observed for Δ^13^C. Our results indicate that host phylogeny, attachment site and host stable isotope value drive Δ^15^N of helminths in coral reef fish while Δ^13^C is more idiosyncratic. These results call for use of taxon- or species-specific and scaled framework for bulk stable isotopes in the trophic ecology of parasites.

## Introduction

Parasitism is the most common life strategy for consumers and is ubiquitous amongst food webs^1^. However, parasites remain a neglected component during the evaluation of biodiversity^2^ and trophic relationships for parasites^3^ remain poorly characterized within food webs as small size, multidisciplinary requirements for identification, and cryptic lifestyles (e.g. multiple hosts associated to multiple larval stages) make identification and characterization of these relationships difficult. The role of parasites in aquatic food chains has been shown to be fundamental^4,5^ and the inclusion of parasitic relationships within food webs dramatically increases the number of trophic links within ecosystems^6^. Despite this, host-parasite relationships remain poorly described, especially for systems with high biodiversity such as coral reefs^7^.

Stable isotope techniques are routinely utilized to study trophic relationships within food webs^8,9^ by using trophic discrimination factors for carbon and nitrogen (Δ^13^C and Δ^15^N) to account for the stepwise increase in δ^13^C and δ^15^N (‰) that occurs between diet and consumer during metabolism^10–12^. However, parasites do not generally follow this relationship with parasite-host discrimination factors (Δ^13^C or Δ^15^N) observed ranging from considerably higher than typical trophic discrimination between predator and prey to negative values for Δ^13^C or Δ^15^N across a variety of taxa^1,13,14^.

Amongst helminths in fish hosts, cestodes and nematodes are usually depleted in δ^15^N values *versus* their hosts^13,15–17^ and vary in Δ^13^C, while trematodes have been found to vary in both Δ^13^C and Δ^15^N^18,19^. However, there can also be considerable variation within these helminth groups^14,20^. Distinct differences for trophic discrimination in parasitic relationships are potentially caused by the combined effects of unique feeding ecologies, often reduced metabolic capabilities of the parasitic taxa being investigated^21,22^, and host metabolic effects due to parasitism^23^. Feeding ecology varies depending on whether the parasite feeds upon host tissue exclusively (on host or within; tissue type dependent^18^), or is able to supplement with material from within the dietary tract as the host feeds or from the environment (e.g. prey items, detritus, mucus, or blood^24^). In addition, it has been suggested that trophic discrimination factors of parasites may not be fixed but scale with the isotopic signature of their hosts, both within^25^ and among parasite species^14^.

Despite multiple investigations, clear discrimination patterns between taxa have not emerged, making simple incorporation of parasites into food web studies using a single universal trophic discrimination factor impossible. This knowledge gap warrants further investigation into the drivers of discrimination factors in helminths. In this study, we examine both δ^13^C and δ^15^N values from whole tissue of 136 helminth parasite-host pairings from coral reef fish to determine the isotopic relationship between taxonomically distinct groups of helminth parasites and to investigate the effect of the habitat in the host and host isotopic signature on helminth isotopic discrimination. We expect that parasite isotopic enrichments and variability *versus* their host will be larger in parasites located in the dietary tract while parasites solely utilizing host tissues will have less variability in their isotopic discrimination. In addition, we expect a negative scaling of parasite discrimination factors with the δ^13^C and δ^15^N values of their hosts.

## Results

We examined the isotopic discrimination of 136 helminth parasite-host pairings including trematodes (n=27), cestodes (n=19), and nematodes (n=90) from 4 host reef lagoon-associated fish species (Table 1, Fig. 1). Trematodes and five of the nematode species were sampled from the dietary tracts (DT) of host species while the remainder were samples from the general cavity (GC). Six of the Δ^15^N values for parasite-host pairs were negative (Supplementary Table 1), with positive relationships (1.05 to 1.58‰) predominately occurring in dietary tract associated parasites (Fig. 1). In 9 cases δ^15^N values were statistically significantly different between parasites and hosts (Supplemental Table 1 & Fig. 2), excluding two pairs from the general cavity: *A. novacaledonica-L. genivitattus* and *Philometra* sp.-*S. undosquamis* (0.58 and 1.18‰, respectively). For the three parasite pairs found in both *L. genivittatus* and *N. furcosus* (i.e. *A. novacaledonica, Callamanus* sp., and *Pseudophyllidae*), Δ^13^C and Δ^15^N were consistently the same between the parasite-host pairings within the same host, either positive or negative, except for carbon in *A. novacaledonica* (0.47‰ *L. genivitattus versus* −1.10‰ *N. furcosus*) (Supplemental Table 1, Fig. 2). Comparison of Δ^15^N between taxa and tissue types for nematodes found significant differences (One-way ANOVA: F_3, 132_: 26.9 *p*< 0.001; Fig. 3). Host δ^15^N was examined *versus* Δ^15^N and found that a linear regression for all of the pairings combined minus the herbivore pairing had a negative slope of −1.2 (R^2^ = 0.071; Fig. 4) with negative slopes observed within individual pairings that were statistically different than 0 for 7 of the parasite-host pairings at α=0.05 (Fig. 4; Supplementary Table 2).

**Figure 1:**
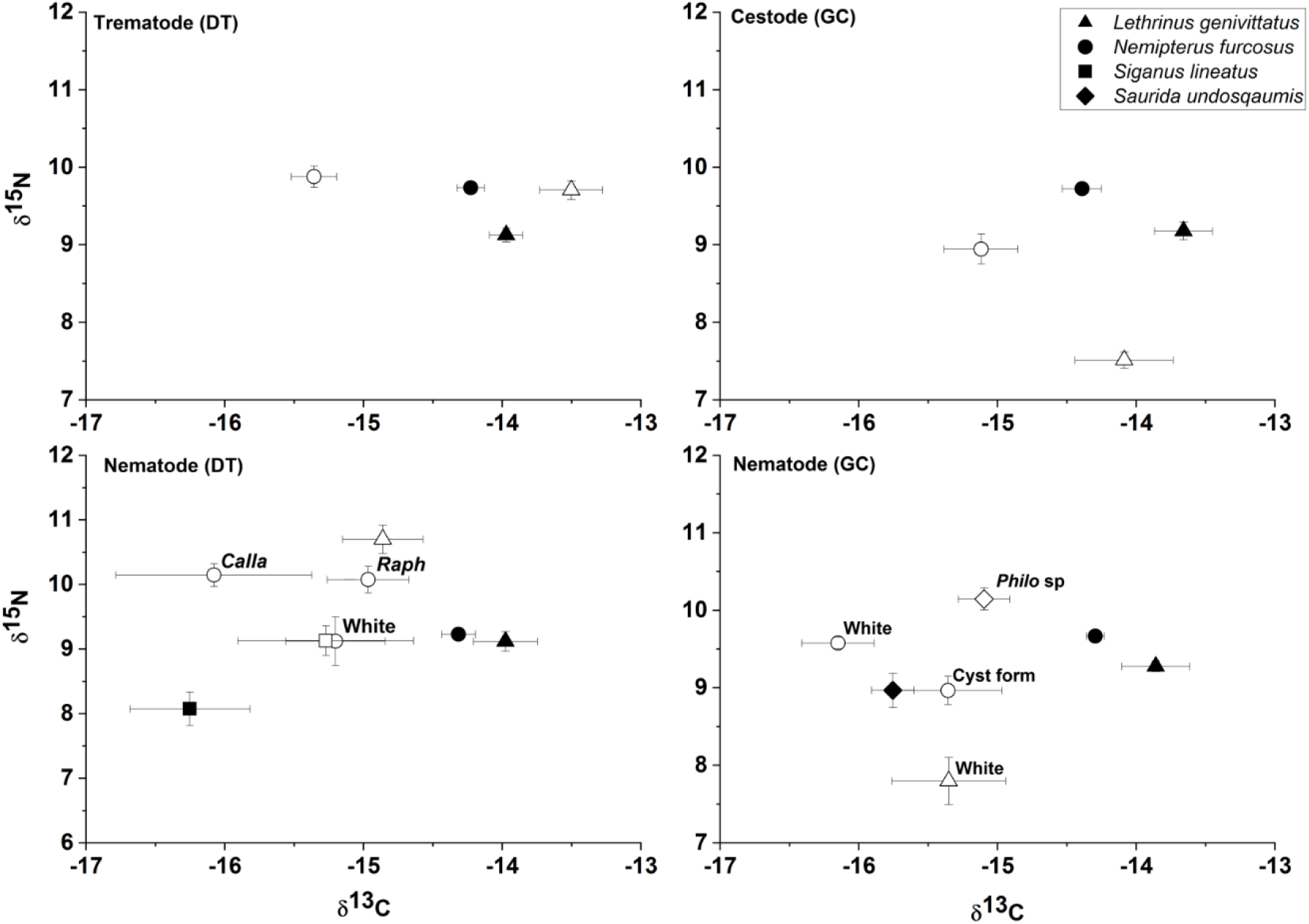
Mean δ^15^N and δ^13^C composition (± SD) for parasite and reef fish host pairings. Parasites and hosts are displayed with open and filled symbols, respectively. Dietary tract (DT) and general cavity (GC) refer to where the parasites were found within fish hosts.

**Figure 2:**
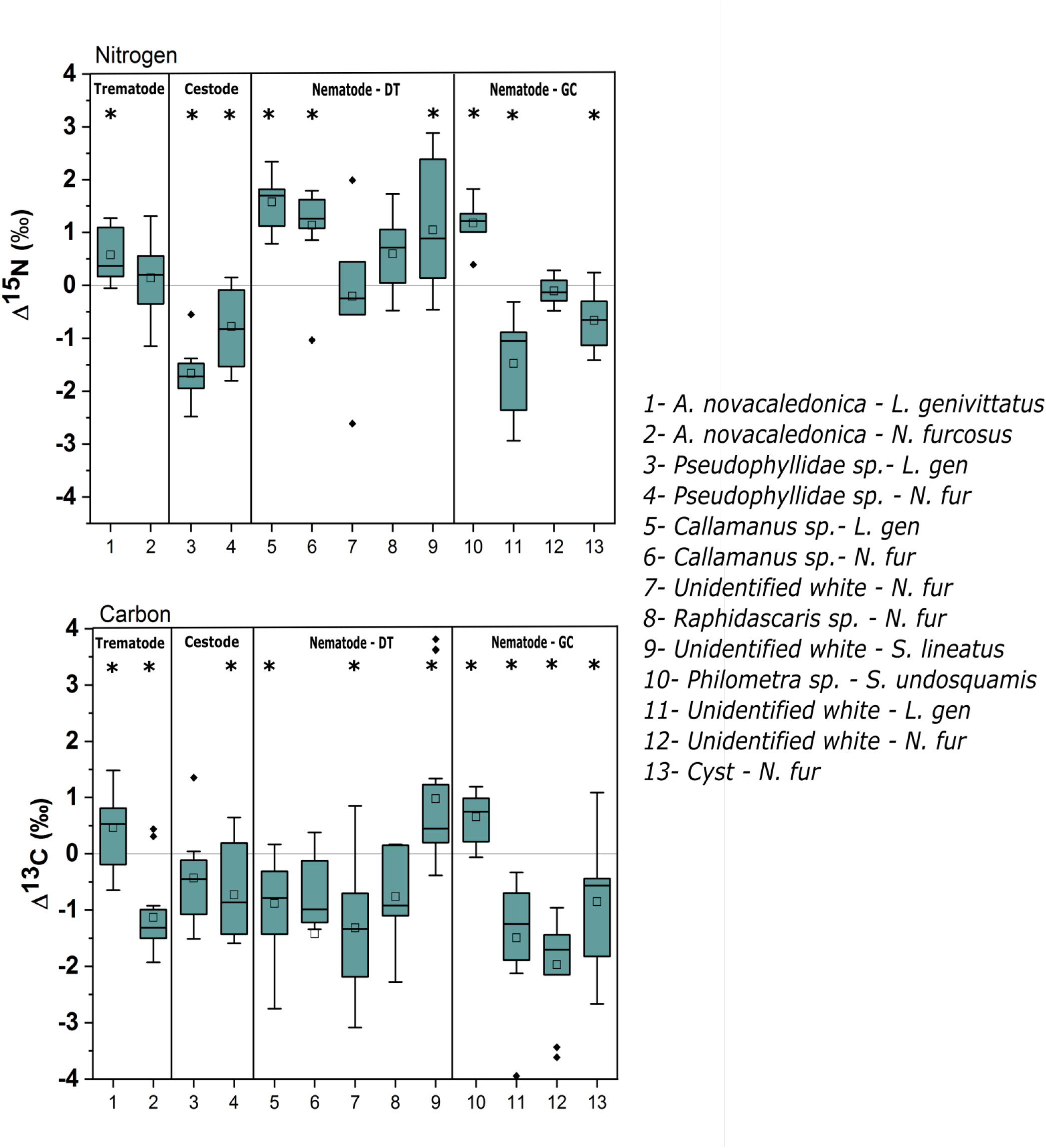
Δ^15^N and Δ^13^C values (‰) for parasite-host couplings examined in this study. Asterisks indicate statistically significant differences between host and parasite at an α = 0.05. Dietary tract (DT) and general cavity (GC) refer to where the parasites were found within fish hosts. For the boxplots, unfilled squares indicate mean, lines indicate median, boxes indicate upper and lower quartiles, and whiskers indicate 1.5 quartile ranges. Black diamonds are outliers.

**Figure 3:**
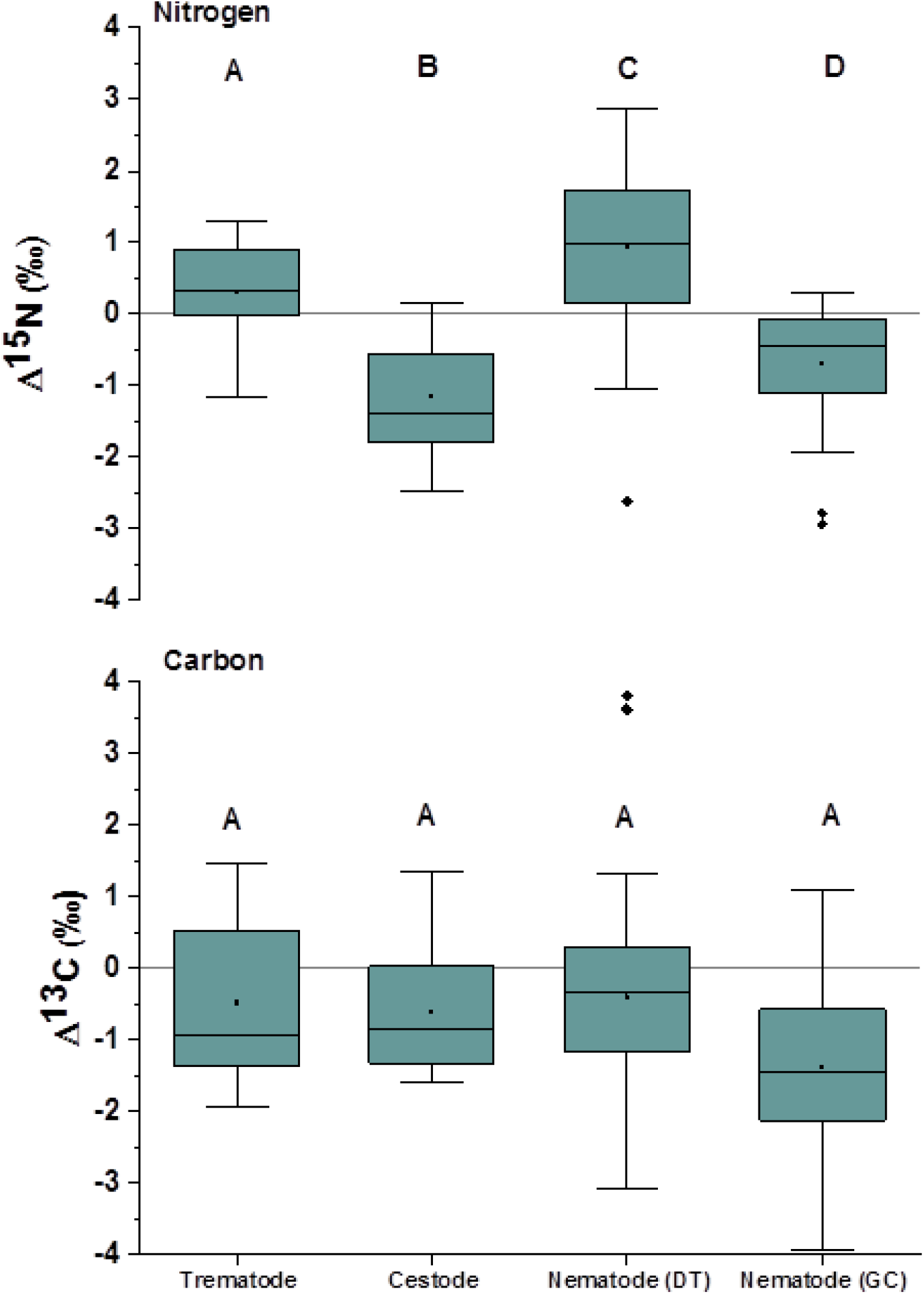
Δ^15^N and Δ^13^C values (‰) for taxa and nematodes separated by infection site. Letters indicate significant differences (*post hoc* Tukey’s test, α=0.05). Dietary tract (DT) and general cavity (GC) refer to where the parasites were found within fish hosts. For the boxplots, black dots indicate mean, lines indicate median, boxes indicate upper and lower quartiles, and whiskers indicate 1.5 quartile ranges. Black diamonds are outliers.

**Figure 4:**
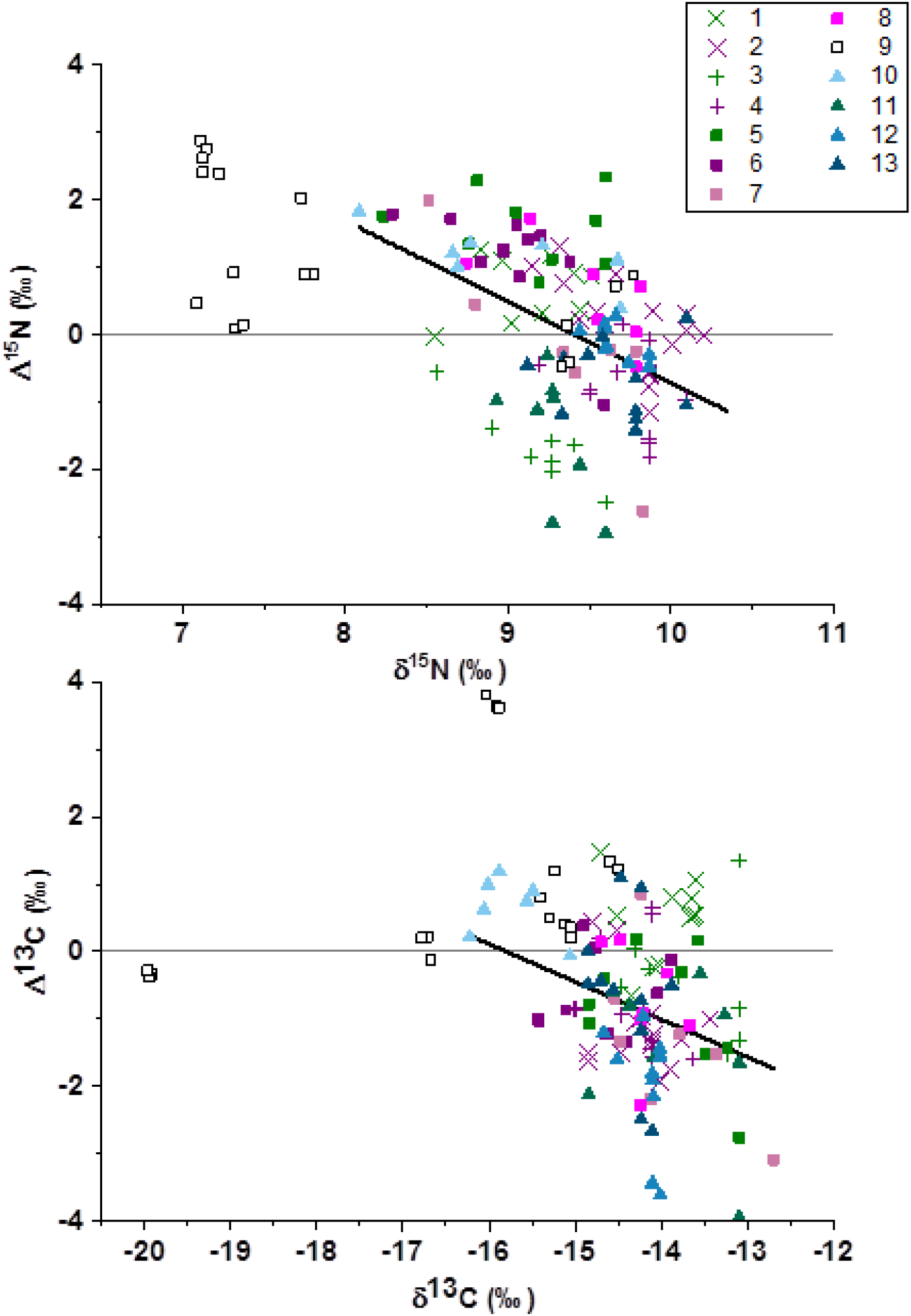
Host δ^15^N values versus Δ^15^N values classified for each individual host-parasite pairing. Labeling follows the pairing order 1-13 from Figure 2 with X indicating trematodes, + indicating cestodes, squares indicating dietary tract nematodes and triangles indicating gut cavity nematodes. Regression lines indicate the statistically significant relationships for the combined pairings from the three predatory fish in the dataset (i.e. excluding samples from the herbivore host *S. lineatus* (open squares, 9)) for nitrogen (slope=-1.2, R^2^ = 0.071, *p* 0.001) and carbon (slope=-0.56, R^2^= 0.196, *p*=0.17).

**Table 1.** Number of parasite-host pairings analysed from different attachment sites, the digestive tract (DT) or the general cavity (GC), from four fish hosts sampled in the coral reef lagoon of New Caledonia.

| Fish host (size range, cm) | Parasite | Attachment site on host (DT / GC) | N |
| --- | --- | --- | --- |
| *Lethrinus genivittatus*<br>(11.0 – 21.5) | *Adlardia novacaledonica* (Trematode) | DT | 11 |
|  | Pseudophyllidae (Cestode) | GC | 8 |
|  | *Callamanus* sp. (Nematode) | DT | 9 |
|  | Unidentified white nematode | GC | 8 |
| *Nemipterus furcosus*<br>(12.1 – 25.0) | *Aldardia novacaledonica* (Trematode) | DT | 16 |
|  | Pseudophyllidae (Cestode) | GC | 11 |
|  | *Callamanus* sp. (Nematode) | DT | 11 |
|  | *Rhaphidascaris* sp. (Nematode) | DT | 7 |
|  | Unidentified white/cream nematode | GC | 10 |
|  | Unidentified nematode (cyst form) | GC | 13 |
|  | Unidentified white nematode | DT | 7 |
| *Siganus undosquamis*<br>(17.8 – 25.6) | *Philometra* sp. (Nematode) | GC | 7 |
| *Saurida lineatus*<br>(28.1 – 38.0) | Unidentified white nematode | DT | 18 |

Δ^13^C values of the pairings were generally negative with lower δ^13^C in the parasite than those of the host fish, except for *S. undosquamis* / *Philometra* sp. (Δ^13^C −0.06 to 1.19‰), *S. lineatus* / white nematode (Δ^13^C −0.38 to 3.81‰) and *L. genivittatus* / *A. novacaledonica* (Δ^13^C 0.47 to 0.98‰, Figs. 2 & 3). The first two of these relationships occur for parasites in the digestive tract and the last one in the general cavity of the host fish. Among the 13 fish-parasite pairings tested, statistically significant differences between host and parasite δ^13^C were observed for 10 cases with a Δ^13^C range from −0.73 to −1.97 ‰. There was no clear Δ^13^C distinction observed between parasites found in the digestive tract *versus* those found in the general cavity. Comparison of Δ^13^C between taxa and tissue types for nematodes found no significant differences (One-way ANOVA: F_3, 132_:1.4 *p*= 0.2; Fig. 3). Host δ^13^C was examined *versus* Δ^13^C and found that a linear regression for all of the pairings combined minus the herbivore pairing had a negative slope of −0.56 (R^2^ =0.196, *p*<0.001; Fig. 4), but that linear regressions for individual pairings indicated no statistically significant difference from 0 for those relationships (Supplementary Table 2).

Mean δ^13^C and δ^15^N values for the host species *L. genivittus* were −14.04 ± 0.51‰ and 8.97 ± 0.52‰, *N. furcosus* were −14.16 ± 0.49‰ and 9.31 ± 0.50‰, *S. undosquamis* were - 16.25 ± 0.49‰ and 8.07 ± 1.09‰, and *S. lineatus* were −15.75 ± 0.40‰ and 8.97 ± 0.59‰, respectively (Fig. 1; Supplementary Table 1), with significant differences observed between species for both δ^13^C and δ^15^N values (One-way ANOVA: δ^13^C, *F*_3,177_= 52.6, *p*< 0.001; δ^15^N, *F*_3,177_=22.8, *p*< 0.001). Body size did not influence δ^13^C and δ^15^N values for *S. undosquamis* and *S. lineatus* for the size range and number of individuals considered here (Pearson correlation, p> 0.05 in all cases). By contrast, fish size significantly influenced δ^15^N values for *L. genivittus* and *N. furcosus* (p< 0.001 for both species; Fig. 5) but not for δ^13^C values (p> 0.05). Calculated trophic levels were similar for the whole populations of *L. genivittus* and *S. undosquamis* (2.86 and 2.79, respectively) with *S. lineatus* having the lowest and *N. furcosus* having the highest trophic levels (2.52 and 2.99, respectively; One-way ANOVA, F_3,137_=42.7, *p*< 0.001). Trophic levels were also higher for larger fish for both *L. genivattus* (2.6 and 2.86 for 11-15 cm and 18-21.5 cm individuals, respectively; one-way ANOVA: F_1,39_, *p*< 0.001) and *N. furcosus* (2.78 and 2.99 for 12.1-15 cm and 20-25.3 cm individuals, respectively; one-way ANOVA: F_1,61_=27.6, *p*< 0.001).

**Figure 5:**
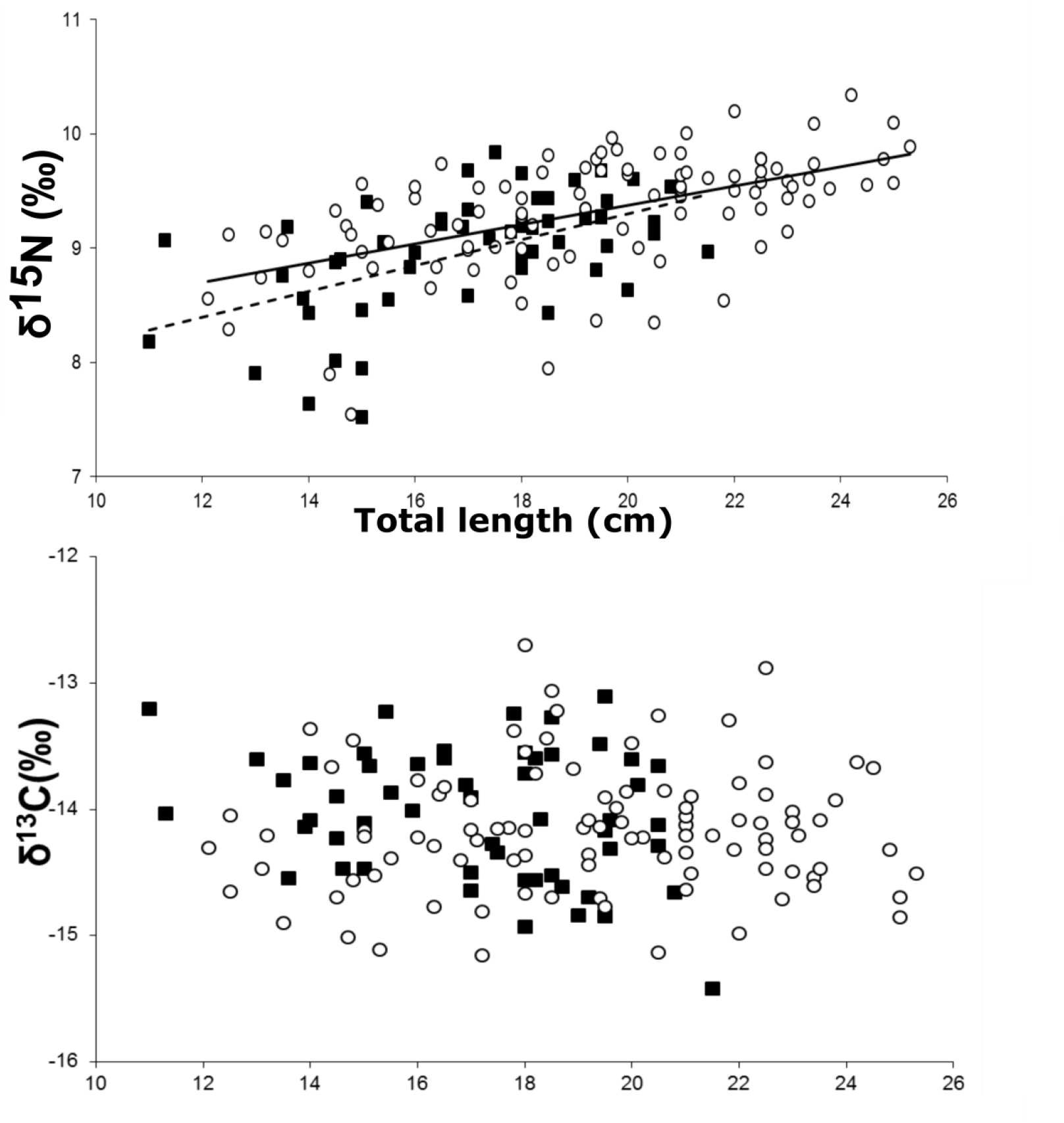
Relationship between fish total length (cm) and δ^15^N (‰) and δ^13^C (‰). Pearson correlations indicated significant increase in δ^15^N values for both *L. genivittatus* (solid squares, dashed line, R^2^=0.30,*p*<0.001) and *N. furcosus* (open circles, solid line, R^2^= 0.31, *p*<0.001) with no corresponding increase for δ^13^C values.

## Discussion

Δ^15^N varied inconsistently within and between taxa, with the most consistent result being elevated Δ^15^N (>0‰) for dietary tract associated nematodes likely associated with feeding on host dietary items in addition to tissue. Δ^13^C was consistently negative between parasite taxa and likely indicates increased reliance on fatty acids from the host to support tissue growth in reef fish-associated helminths. The varied relationships amongst and between taxa provide further evidence that parasite-host pairings are distinctly different than typical trophic relationships and warrant further investigation to adequately characterize parasite contributions to food webs.

Δ^15^N values showed no difference or were positive for the dietary tract associated trematode and nematodes *(Allardia, Callamanus, Rhaphidascaris*, and unidentified white) while the cestodes had negative values for both pairings examined. Δ^15^N values for the general cavity-associated nematodes varied, with a strong positive value for the gonad-associated Philometra sp.- *S. undosquamis* pairing, no difference observed for unidentified white nematode - *N. furcosus* pairing, and strong negative relationships for both the nematode cyst-type – *N. furcosus* and the unidentified white nematode – *L. genivittatus* pairings (Fig. 2).

Trematodes have been found to have positive to neutral Δ^15^N values regardless of infection site (e.g. dietary tract or general cavity^18,19^) indicating utilization of host-metabolized nitrogen derived from tissue in addition to nitrogen compounds derived from the host diet. The low Δ^15^N values (0.18 to 0.6‰) observed in this study demonstrate close association of the trematode with the host diet. Sole reliance on host tissue and therefore host-metabolized N would be expected to yield a “classical” trophic enrichment of ~3.4‰, which is considerably larger than the observed Δ^15^N. Small Δ^15^N values likely reflect combined utilization of more ^15^N enriched sources of nitrogen derived from host tissue as well as more ^15^N depleted compounds either metabolized or directly assimilated. Negative Δ^15^N values have been previously observed for cestodes within fish hosts^15,16,18,19,24,26^ and may be caused by direct utilization of relatively ^15^N-depleted compounds from the host diet^16^ or metabolically recycled ^15^N-depleted AAs produced by the gut microbial community^27^ that both cestodes and trematodes are well-positioned to utilize while residing in the dietary tract.

Similarly, three of the general-cavity associated nematodes displayed negative or neutral Δ^15^N values, indicating that direct uptake from the host without further metabolic processing by the parasite of nitrogen compounds as a likely pathway for N in this taxa. This uptake was not consistent across the taxa, with different species that target similar infection sites (e.g. dietary tract *versus* general cavity) displaying considerably different Δ^15^N values. The single gonad-associated nematode (*Philometra* sp. - *S. undosquamis*) examined had a positive Δ^15^N indicating at least partial reliance on direct utilization of metabolized N from host tissues. The dietary tract-associated nematodes were generally ^15^N-enriched in comparison to their hosts indicating at least partial utilization of host-metabolized compounds from tissue or the taxa-dependent ability for nematodes to biosynthesize AAs from nitrogenous compounds^22^.

Δ^13^C values were predominately neutral or negative for ten of the pairings examined, with positive Δ^13^C only observed for the *L. genivittatus - A. novacaledonica*, unidentified white – *S. lineatus*, and *Philometra* sp. - *S. undosquamis* pairings. No change or depletion in δ^13^C does not agree with the expected 0.5-1‰ increase that is usually expected for trophic interactions^11^, but likely reflects reliance on lipids and fatty acids directly derived from the host or host diet to support helminth tissue growth. Plathelminthes and some nematodes have been found to be incapable of *de novo* fatty acid synthesis and to have to rely on fatty acids derived from the host^22,28^ due to incomplete metabolic pathways for lipid biosynthesis. Direct uptake of fatty acids and other lipids from the host would be expected to coincide with a minimum carbon fractionation as no further metabolic processing is required, thereby maintaining the relatively low δ^13^C values associated with lipids as they are incorporated into the parasite. The relatively uniform neutral or negative relationships for Δ^13^C values across the pairings located from both the dietary tract and the general cavity indicate that lipid carbon is likely utilized to support tissue growth beyond species closely associated with fatty tissues (e.g. blood and liver)^13,14^. This relationship should be examined further with methods that incorporate metabolic pathway techniques for target species beyond model organisms targeted for pathogenicity^21,22^. Pairings that have elevated Δ^13^C values likely reflect decreased utilization of host lipids and increased reliance on either host sugars or proteins processed through more complete metabolic pathways within the helminths to provide tissue carbon or potentially different δ^13^C compositions from different host tissues^18^.

δ^15^N values for both *L. genivattus* and *N. furcosus* increased with body size leading to higher trophic levels in larger fish, a pattern commonly observed for coral reef-associated fish, while there was no corresponding increase or shift in δ^13^C throughout ontogeny. No change in δ^13^C indicates that both smaller and larger individuals are relying on similar sources of underlying carbon production^29^, and that the larger individuals are feeding on larger prey with an elevated trophic level, i.e. elevated δ^15^N values^30^.

Additionally, negative scaling of both Δ^13^C and Δ^15^N values *versus* δ^13^C and δ^15^N of fish hosts were observed with statistically significant negative slopes observed within pairings for nitrogen in 7 of the 13 parasite-host pairs (Fig. 4), while no statistically significant within-pairings relationships were found for carbon (Supplementary Table 2). Negative scaling of trophic discrimination factors with increased host carbon and nitrogen values has previously been observed for parasite-host, predator-prey and herbivore-plant relationships^12,14,31^ and may be associated with dietary quality. In this study, there appears to be an increased spread in the fractionation (Δ^15^N values) observed within the herbivore *S. lineatus*, with the lowest δ^15^N values occurring with the highest Δ^15^N values and a distinct grouping of individuals with lower Δ^15^N values corresponding with the highest δ^15^N values (Fig. 4). This wide range of values may represent the relative richness in diet, with strictly herbivorous individuals causing a shift in their parasites towards exclusive utilization of host tissues, while individuals that supplement with animal protein (more omnivorous) provide additional material within their diet for their parasites to supplement from. Increased protein quality in a predator’s diet results in a smaller ‰ difference between the diet and consumer, i.e. a smaller trophic fractionation^32^. This relationship coincides with the trend of decreased Δ^15^N values being observed for increased trophic level predation within predators in this study (Fig. 4). In herbivores, larger trophic discrimination factors for nitrogen are often observed^33^, and supplementation with protein (omnivory) would be expected to generate a negative offset in Δ^15^N if the parasites are supplementing from dietary protein in addition to host tissues.

In conclusion, this study characterized discrimination factors for carbon and nitrogen within helminths living in coral reef fish and highlights the uncertainties that remain in adequately describing parasitic relationships within food webs. These uncertainties call for the development of a taxon- or species-specific and scaled framework for using bulk stable isotope analysis to study the trophic ecology of parasites. In addition, further work using metabolomics and compound specific stable isotope techniques is warranted in order to better characterize the underlying metabolic differences that are driving the differences observed for trophic discrimination factors between parasite and hosts.

## Methods

### Sampled areas and studied species

Individual fish were captured in New Caledonia, southwestern Pacific Ocean. The three species *Lethrinus genivittatus* Cuvier & Valenciennes, 1830, *Nemipterus furcosus* (Cuvier & Valenciennes, 1830) and *Saurida undosquamis* (Richardson, 1848) were caught using hand lines in the lagoon off the city of Nouméa (22°18’S and 166°25’E) at approximately 10-12 m depth, in August 2011, 2013 and 2014. Three years of data from catches were pooled as a preliminary two-way ANOVA (year x size) revealed that year was not a significant factor (p > 0.05). *L. genivittatus* feeds on crabs and worms, *N. furcosus* feeds on crabs and shrimp and *S. undosquamis* is predominantly piscivorous^34^. The species *Siganus lineatus* (Cuvier & Valenciennes, 1835) was caught in coastal mangroves in the southeast coast at Yaté (22°16’S and 167°0l’l5E) using gillnets in June-August 2014. This species is usually considered an herbivore^35^, but has been observed to predominately feed on algae and supplement with minor consumption of invertebrates in Yaté^36^. Parasites were present in all fish that were examined and appear to be ubiquitous within the species examined in this study.

All individuals caught were immediately placed in ice until further processing in the laboratory. Each fish was measured to the nearest 0.1 cm (total length) and a small piece of dorsal muscle of each fish was sampled and immediately frozen at −20°C. To extract the parasites, the general cavity was first examined to collect parasites found outside the digestive tracts embedded in or attached to fish tissues. In a second step, the method presented in Justine, et al. was applied to extract the parasites found alive within the digestive tracts. All parasites having a sufficient biomass were collected and immediately frozen, i.e. nematodes, cestodes and trematodes. A total of 54 *L. genivittatus* were caught with 36 exploitable fish-parasite pairings, 99 *N. furcosus* with 75 exploitable fish-parasite pairings, and respectively 7 *S. undosquamis* and 18 *S. lineatus* were exploitable as fish-parasite pairings (Table 1).

### Stable isotope preparation and analyses

Carbon and nitrogen stable-isotope ratios (δ^13^C and δ^15^N) were determined for dorsal muscle tissue of all fishes collected. Fish muscle tissue is routinely utilized for stable isotope values for fish as it does not require lipid extraction prior to analysis^38^. Samples were freeze-dried and ground into a fine powder using a mortar and pestle. One milligram of powdered material was loaded into tin capsules and analysed for each sample without prior treatment. This same procedure was used for parasites (whole animal) for samples that had sufficient dry mass (≥ 0.3 mg).

^13^C/^12^C and ^15^N/^14^N ratios were determined with a continuous-flow isotope-ratio mass spectrometer (Thermo Scientific Delta V Advantage, Bremen, Germany) coupled to an elemental analyser (Thermo Scientific Flash EA1112, Bremen, Germany). The analytical precision was 0.1‰ for ^13^C and 0.15‰ for ^15^N, estimated using the internal standards leucine calibrated against ‘Europa flour’ and IAEA standards N1 and N2. Isotope ratios were expressed as δ notation (‰) differences from a standard reference material:

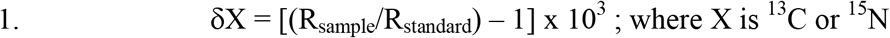

Where R is the corresponding ratio (^13^C/^12^C or ^15^N/^14^N) for both sample and reference standard and δX is the measured isotopic value in per mil (‰) relative to the international standard references are Vienna Pee Dee Belemnite (vPDB) for carbon and atmospheric N_2_ for nitrogen.

Parasite-host discrimination factors were calculated using:

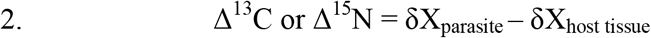

Where δX represents the isotopic value of carbon or nitrogen for each parasite-host tissue pairing examined.

### Data analysis

The significance of differences in δ^13^C and δ^15^N between a fish and its parasite was tested with the Wilcoxon signed rank test when homogeneity of variances was not verified or paired samples t-tests when homogeneity of variances was verified, dependant on fish species. The relationships between fish size and isotopic values (δ^15^N or δ^13^C) were investigated with Pearson correlation coefficients. One-way analysis of variance (ANOVA) was used to determine significant differences between host and parasite δ^13^C and δ^15^N values and to explore the relationship between host size and trophic level. The relationship between host δ^13^C and δ^15^N values and Δ^13^C and Δ^15^N values were determined through linear regression followed with subsequent application of an F test of the modelled slope against a slope of 0. This was done among and within the host-parasite parings. For the analysis among pairings we excluded samples from the herbivorous fish host *S. lineatus* (19 samples) due to their very different isotope values to avoid skewing the relationship due to explained outliers. The trophic level (TL) of fish individuals was calculated following the formulae of^10^:

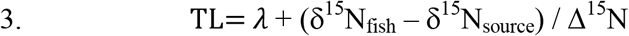

where *λ* is the trophic level of the source of organic matter, i.e. 1,: δ^15^N_fish_ is the isotopic value of nitrogen for the considered fish, δ^15^N_source_ is the isotopic value of the source of organic matter at the base of the food web, i.e. 3.59 for sedimentary organic matter^29^ that concerns *L. genivittatus, N. furcosus* and *S. undosquamis*, all caught of sandy unvegetated bottom; and 2.12 for the most eaten algae by *Siganus lineatus* and Δ^15^N that is the trophic enrichment factor (TEF) between a food item and its consumer. Here, we adopted a value of 3.9 ‰ for *S. lineatus*^36,39^ reflecting usually higher TEF for herbivores compared to the conventional 3.4 ‰ value^33^. For the three other species, we adopted a TEF of 3.0 ‰ because TEF are usually lower than the conventional value for carnivores^33,40^.

## Acknowledgments

We are grateful to the students in “Licence Sciences de la Vie, de la Terre et de l’Environnement” de l’Université de la Nouvelle-Calédonie for their valuable help for the parasites sampling during dissections, to C. Pigot for his help during fish catches and samples preparation, to G. Guillou for stable isotopes analyses.

## Additional Information

Declarations of interest: none. MB, TM, YL and PS contributed to experimental design, sample collection and processing, and data collation. PMR wrote the manuscript with considerable consultation from DT, MvdM and YL for data analysis. All authors reviewed the manuscript prior to submission.

## References

1 Lafferty, K. D. et al. Parasites in food webs: the ultimate missing links. Ecology Letters 11, 533–546, doi:10.1111/j.1461-0248.2008.01174.x (2008).

2 Poulin, R. Parasite species richness in New Zealand fishes: a grossly underestimated component of biodiversity? Diversity and Distributions 10, 31–37 (2004).

3 Lafferty, K. D. & Kuris, A. M. Trophic strategies, animal diversity and body size. Trends in Ecology & Evolution 17, 507–513 (2002).

4 Welicky, R. L., Demopoulos, A. W. J. & Sikkel, P. C. Host-dependent differences in resource use associated with *Anilocra* spp. parasitism in two coral reef fishes, as revealed by stable carbon and nitrogen isotope analyses. Marine Ecology 38, doi:10.1111/maec.12413 (2017).

5 Kuris, A. M. et al. Ecosystem energetic implications of parasite and free-living biomass in three estuaries. Nature 454, 515–518, doi:10.1038/nature06970 (2008).

6 Dunne, J. A. et al. Parasites affect food web structure primarily through increased diversity and complexity. PLoS biology 11 (2013).

7 Justine, J.-L. Parasite biodiversity in a coral reef fish: twelve species of monogeneans on the gills of the grouper Epinephelus maculatus (Perciformes: *Serranidae*) off New Caledonia, with a description of eight new species of *Pseudorhabdosynochus* (Monogenea: *Diplectanidae’)*. Systematic Parasitology 66, 81 (2007).

8 Minagawa, M. & Wada, E. Stepwise enrichment of ^15^N along food chains: Further evidence and the relation between δ^15^N and animal age. Geochimica et Cosmochimica Acta 48, 1135–1140, doi:https://doi.org/10.1016/0016-7037(84)90204-7 (1984).

9 Fry, B. Food web structure on Georges Bank from stable C, N, and S isotopic compositions. Limnology and Oceanography 33, 1182–1190, doi:10.4319/lo.1988.33.5.1182 (1988).

10 Post, D. M. Using stable isotopes to estimate trophic position: models, methods, and assumptions. Ecology 83, 703–718, doi:10.1890/0012-9658(2002)083[0703:usitet]2.0.co;2 (2002).

11 McCutchan, J. H., Lewis Jr, W. M., Kendall, C. & McGrath, C. C. Variation in trophic shift for stable isotope ratios of carbon, nitrogen, and sulfur. Oikos 102, 378–390, doi:10.1034/j.1600-0706.2003.12098.x (2003).

12 Caut, S., Angulo, E. & Courchamp, F. Variation in discrimination factors (Δ^15^N and Δ^13^C): the effect of diet isotopic values and applications for diet reconstruction. Journal of Applied Ecology 46, 443–453, doi:10.1111/j.1365-2664.2009.01620.x (2009).

13 Pinnegar, J. K., Campbell, N. & Polunin, N. V. C. Unusual stable isotope fractionation patterns observed for fish host—parasite trophic relationships. Journal of Fish Biology 59, 494–503, doi:10.1111/j.1095-8649.2001.tb02355.x (2001).

14 Thieltges, D. W., Goedknegt, M. A., O’Dwyer, K., Senior, A. M. & Kamiya, T. Parasites and stable isotopes: a comparative analysis of isotopic discrimination in parasitic trophic interactions. Oikos 128, 1329–1339, doi:10.1111/oik.06086 (2019).

15 Deudero, S., Pinnegar, J. K. & Polunin, N. V. C. Insights into fish host-parasite trophic relationships revealed by stable isotope analysis. Diseases of Aquatic Organisms 52, 77–86 (2002).

16 Power, M. & Klein, G. M. Fish host-cestode parasite stable isotope enrichment patterns in marine, estuarine and freshwater fishes from northern Canada. Isotopes in environmental and health studies 40, 257–266 (2004).

17 Navarro, J. et al. Isotopic discrimination of stable isotopes of nitrogen (δ^15^N) and carbon (δ^13^C) in a host-specific holocephalan tapeworm. Journal of helminthology 88, 371–375 (2014).

18 Kamiya, E., Urabe, M. & Okuda, N. Does atypical ^15^N and ^13^C enrichment in parasites result from isotope ratio variation of host tissues they are infected? Limnology, doi:10.1007/s10201-019-00596-w (2019).

19 Kanaya, G. et al. Application of stable isotopic analyses for fish host-parasite systems: an evaluation tool for parasite-mediated material flow in aquatic ecosystems. Aquatic Ecology 53, 217–232, doi:10.1007/s10452-019-09684-6 (2019).

20 Gilbert, B. M. et al. You are how you eat: differences in trophic position of two parasite species infecting a single host according to stable isotopes. Parasitology Research, 1–8 (2020).

21 International Helminth Genomes, C. Comparative genomics of the major parasitic worms. Nat Genet 51, 163–174, doi:10.1038/s41588-018-0262-1 (2019).

22 Tyagi, R., Rosa, B. A., Lewis, W. G. & Mitreva, M. Pan-phylum Comparison of nematode metabolic potential. PLoS Negl Trop Dis 9, e0003788–e0003788, doi:10.1371/journal.pntd.0003788 (2015).

23 Yohannes, E., Grimm, C., Rothhaupt, K.-O. & Behrmann-Godel, J. The effect of parasite infection on stable isotope turnover rates of δ^15^N, δ^13^C and δ^34^S in multiple tissues of eurasian perch *Perca fluviatilis*. PLOS ONE 12, e0169058, doi:10.1371/journal.pone.0169058 (2017).

24 Behrmann-Godel, J. & Yohannes, E. Multiple isotope analyses of the pike tapeworm *Triaenophorus nodulosus* reveal peculiarities in consumer–diet discrimination patterns. Journal of Helminthology 89, 238–243, doi:10.1017/S0022149X13000849 (2015).

25 Goedknegt, M. A. et al. Trophic relationship between the invasive parasitic copepod *Mytilicola orientalis* and its native blue mussel (*Mytilus edulis*) host. Parasitology 145, 814–821, doi:10.1017/S0031182017001779 (2017).

26 Persson, M. E., Larsson, P. & Stenroth, P. Fractionation of δ^15^N and δ^13^C for Atlantic salmon and its intestinal cestode *Eubothrium crassum*. Journal of Fish Biology 71, 441–452, doi:10.1111/j.1095-8649.2007.01500.x (2007).

27 McMahon, K. W. & McCarthy, M. D. Embracing variability in amino acid δ^15^N fractionation: mechanisms, implications, and applications for trophic ecology. Ecosphere 7, e01511-n/a, doi:10.1002/ecs2.1511 (2016).

28 Brouwers, J. F., Smeenk, I. M., van Golde, L. M. & Tielens, A. G. The incorporation, modification and turnover of fatty acids in adult *Schistosoma mansoni*. Molecular and biochemical parasitology 88, 175–185 (1997).

29 Briand, M. J., Bonnet, X., Goiran, C., Guillou, G. & Letourneur, Y. Major sources of organic matter in a complex coral reef lagoon: identification from isotopic signatures (δ^13^C and δ^15^N). PloS one 10, e0131555–e0131555, doi:10.1371/journal.pone.0131555 (2015).

30 Greenwood, N. D. W., Sweeting, C. J. & Polunin, N. V. C. Elucidating the 15 13 trophodynamics of four coral reef fishes of the Solomon Islands using δ^15^N and δ^13^C. Coral Reefs 29, 785–792, doi:10.1007/s00338-010-0626-1 (2010).

31 Hussey, N. E. et al. Rescaling the trophic structure of marine food webs. Ecology letters 17, 239–250 (2014).

32 McMahon, K. W., Thorrold, S. R., Elsdon, T. S. & McCarthy, M. D. Trophic discrimination of nitrogen stable isotopes in amino acids varies with diet quality in a marine fish. Limnology and Oceanography 60, 1076–1087, doi:10.1002/lno.10081 (2015).

33 Mill, A. C., Pinnegar, J. K. & Polunin, N. V. C. Explaining isotope trophic-step fractionation: why herbivorous fish are different. Functional Ecology 21, 1137–1145, doi:10.1111/j.1365-2435.2007.01330.x (2007).

34 Kulbicki, M., Guillemot, N. & Amand, M. A general approach to length-weight relationships for New Caledonian lagoon fishes. Cybium 29, 235–252 (2005).

35 Woodland, D. J. Revision of the fish family *Siganidae* with descriptions of two new species and comments on distribution and biology. Indo-Pacific Fishes 19 (1990).

36 Moleana, T. Etude de la reproduction, de l’alimentation et de la composition en acides gras du picot rayé *Siganus lineatus*. Application à la domestication d’une nouvelle espèce tropicale pour la piscuculture marine (Nouvelle-Calédonie; Aqualagon SARL J, Nouvelle Calédonie, (2016).

37 Justine, J.-L., Briand, M. J. & Bray, R. A. A quick and simple method, usable in the field, for collecting parasites in suitable condition for both morphological and molecular studies. Parasitology Research 111, 341–351 (2012).

38 Pinnegar, J. K. & Polunin, N. V. C. Differential fractionation of δ^13^C and δ^15^N among fish tissues: implications for the study of trophic interactions. Functional Ecology 13, 225–231, doi:10.1046/j.1365-2435.1999.00301.x (1999).

39 Abrantes, K. & Sheaves, M. Incorporation of terrestrial wetland material into aquatic food webs in a tropical estuarine wetland. Estuarine, Coastal and Shelf Science 80, 401–412 (2008).

40 Briand, M. J., Bonnet, X., Guillou, G. & Letourneur, Y. Complex food webs in highly diversified coral reefs: Insights from δ^13^C and δ^15^N stable isotopes. Food Webs 8, 12–22, doi:https://doi.org/10.1016/j.fooweb.2016.07.002 (2016).

