## Supplemental tables 1 and 2 for "Impacts of host phylogeny, feeding styles, and parasite attachment site on isotopic discrimination in helminths infecting coral reef fish hosts"

Supplementary Material:

**Supplementary Table 1**. Mean δ**^13^**C and δ**^15^**N signatures (± sd) of fish and parasites, mean Δ**^13^**C and Δ**^15^**N (± sd) for the parasite-host pairings, and associated *p*-value (test for paired samples, see text). Differences in bold characters are statistically significant.

| **Fish host/ parasite pairing** | **δ^13^C** | | Δ**^13^**C | *p* | **δ^15^N** | | Δ**^15^**N | *p* |
| --- | --- | --- | --- | --- | --- | --- | --- | --- |
|  | Fish | Parasite |  |  | Fish | Parasite |  |  |
| *L. genivittatus /A. novacaledonica* | -13.97 (0.40) | -13.50 (0.75) | **0.47 (0.61)** | **0.029** | 9.13 (0.29) | 9.70 (0.40) | **0.58 (0.50)** | **0.003** |
| *L. genivittatus /* Pseudophyllidae | -13.66 (0.59) | -14.09 (1.00) | -0.43 (0.89) | 0.216 | 9.18 (0.32) | 7.51 (0.30) | **-1.66 (0.56)** | **<0.001** |
| *L. genivittatus / Callamanus* sp. | -13.98 (0.69) | -14.86 (0.87) | **-0.88 (0.94)** | **0.022** | 9.12 (0.46) | 10.70 (0.66) | **1.58 (0.54)** | **<0.001** |
| *L. genivittatus /* Unidentified white nematode | -13.86 (0.69) | -15.35 (1.16) | **-1.49 (1.16)** | **0.008** | 9.28 (0.19) | 7.80 (0.86) | **-1.48 (0.96)** | **0.003** |
| *N. furcosus / A. novacaledonica* | -14.22 (0.41) | -15.32 (0.63) | **-1.10 (0.63)** | **<0.001** | 9.76 (0.33) | 9.94 (0.55) | 0.18 (0.68) | 0.280 |
| *N. furcosus /* Pseudophyllidae | -14.39 (0.47) | -15.12 (0.88) | **-0.73 (0.82)** | **0.015** | 9.73 (0.25) | 8.95 (0.64) | **-0.78 (0.66)** | **0.003** |
| *N. furcosus / Callamanus* sp. | -14.66 (0.54) | -16.08 (2.04) | -1.42 (1.47) | 0.085 | 9.01 (0.35) | 10.15 (0.59) | **1.13 (0.78)** | **<0.001** |
| *N. furcosus / Rhaphidascaris* sp. | -14.21 (0.34) | -14.97 (0.78) | -0.76 (0.86) | 0.057 | 9.48 (0.40) | 10.07 (0.54) | 0.60 (0.73) | 0.074 |
| *N. furcosus /* Unidentified white/cream nematode | -13.86 (0.99) | -15.20 (0.50) | **-1.31 (1.23)** | **0.029** | 9.33 (0.50) | 9.12 (0.99) | -0.21 (1.36) | 0.702 |
| *N. furcosus /* Unidentified nematode (cysts) | -14.40 (0.32) | -15.36 (1.29) | **-0.96 (1.07)** | **0.031** | 9.65 (0.31) | 8.97 (0.61) | **-0.69 (0.55)** | **0.002** |
| *N. furcosus /* Unidentified white nematode | -14.18 (0.23) | -16.15 (0.82) | **-1.97 (0.89)** | **<0.001** | 9.68 (0.15) | 9.58 (0.24) | -0.10 (0.25) | 0.227 |
| *S. undosquamis / Philometra* sp. | -15.75 (0.41) | -15.10 (0.49) | **0.66 (0.44)** | **0.008** | 8.97 (0.59) | 10.15 (0.38) | **1.18 (0.43)** | **<0.001** |
| *S. lineatus /* Unidentified white nematode | -16.25 (1.54) | -15.27 (1.39) | **0.98 (1.06)** | **0.007** | 8.08 (1.10) | 9.13 (0.98) | **1.05 (1.07)** | **0.001** |

**Supplementary Table 2**: The slopes from linear regressions of hostδvalues versusΔvalues for the parasite-host pairings for both carbon and nitrogen and the associated R^2^ and *p* values. Bold lettering indicates statistically significant differences.

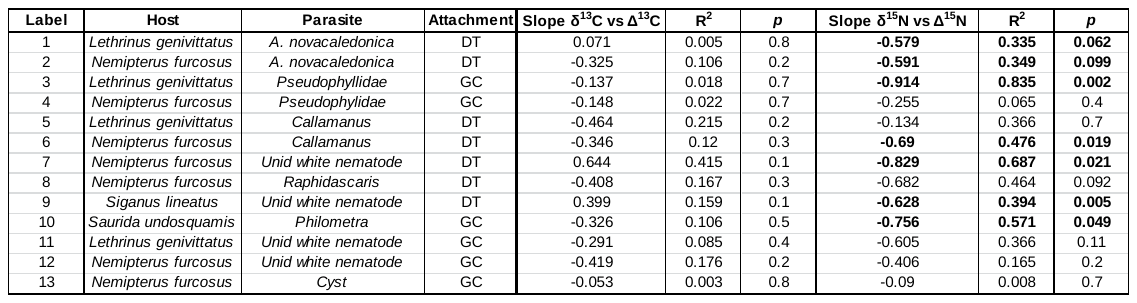
